# Chromosomal distribution of cyto-nuclear genes in a dioecious plant with sex chromosomes

**DOI:** 10.1101/007047

**Authors:** Josh Hough, J. Arvid Ågren, Spencer C. H. Barrett, Stephen I. Wright

**Affiliations:** Department of Ecology and Evolutionary Biology, University of Toronto, Toronto, ON, Canada, M5S 3B2

**Keywords:** Cyto-nuclear interactions, co-adaptation, sexual conflict, gene transfer

## Abstract

The coordination between nuclear and organellar genes is essential to many aspects of eukaryotic life, including basic metabolism, energy production, and ultimately, organismal fitness. Whereas nuclear genes are bi-parentally inherited, mitochondrial and chloroplast genes are almost exclusively maternally inherited, and this asymmetry may lead to a bias in the chromosomal distribution of nuclear genes whose products act in the mitochondria or chloroplasts. In particular, because X-linked genes have a higher probability of co-transmission with organellar genes (2/3) compared to autosomal genes (1/2), selection for co-adaptation has been predicted to lead to an over-representation of nuclear-mitochondrial and nuclear-chloroplast genes on the X chromosome relative to autosomes. In contrast, the occurrence of sexually antagonistic organellar mutations might lead to selection for movement of cyto-nuclear genes from the X chromosome to autosomes to reduce male mutation load. Recent broad-scale comparative studies of N-mt distributions in animals have found evidence for these hypotheses in some species, but not others. Here, we use transcriptome sequences to conduct the first study of the chromosomal distribution of cyto-nuclear interacting genes in a plant species with sex chromosomes (*Rumex hastatulus*; Polygonaceae). We found no evidence of under- or over-representation of either N-mt or N-cp genes on the X chromosome, and thus no support for either the co-adaptation or the sexual-conflict hypothesis. We discuss how our results from a species with recently evolved sex chromosomes fit into an emerging picture of the evolutionary forces governing the chromosomal distribution of nuclear-mitochondrial and nuclear-chloroplast genes.

## Introduction

The intimate relationships between nuclear and organellar genomes in eukaryotes represent some of the most striking examples of co-evolved mutualisms (Gillham 1994; Lane 2005; Aanen et al. 2014). The long co-evolutionary history of nuclear and mitochondrial genomes is perhaps best illustrated by the finding that the vast majority of mitochondrial genes in animals have been transferred to the nuclear genome (Adams and Palmer 2003; Rand et al. 2004; Burt and Trivers 2006). Indeed, animal mitochondria now encode only a few proteins after having lost the majority of their original genes (Berg and Kurland 2000; Ridley 2000; Bar-Yaacov et al. 2012). Moreover, almost one fifth of the *Arabidopsis thaliana* nuclear genome is of chloroplast origin (Martin 2003), suggesting that organellar-to-nuclear gene movement has played a crucial role in the evolution of plant genetic systems.

The evolution of cyto-nuclear interactions and the chromosomal distribution of the genes involved should be influenced by the contrasting modes of inheritance of organellar genes (maternal inheritance) and autosomal genes (bi-parental inheritance). This difference may, for example, result in conflict between nuclear and organellar genes over sex determination and sex ratio (Cosmides and Tooby 1981; Werren and Beukeboom 1998), and several mitochondrial genes in plants are known to cause male sterility (Burt and Trivers 2006; Touzet and Meyer 2014). In systems with XY sex determination, where males are the heterogametic (XY) and females the homogametic sex (XX), genes on the X chromosome spend 2/3 of their time in females (Rand et al. 2001) and therefore share a female-biased inheritance pattern relative to Y-linked or autosomal genes, which may result in inter-genomic co-adaptation or conflict.

A potential consequence of inter-genomic conflict or co-adaptation between nuclear genes, whose products interact with mitochondrial or chloroplasts (mito-nuclear and cyto-nuclear genes, respectively) and other regions of the genome, is a shift in the chromosomal location of such genes, either becoming more or less abundant on the X chromosome. Several molecular mechanisms have been suggested to be involved in driving gene movement, including gene duplication followed by fixation and subsequent gene loss (Wu and Yujun Xu 2003), and autosomal gene duplications followed by the evolution of sex biased gene expression (Connallon and Clark 2011). The evolutionary mechanisms of this gene movement have also been explored by several recent studies (Drown et al. 2012; Hill and Johnson 2013; Dean et al. 2014; Rogell et al. 2014), and two main processes have been proposed to account for the movement of genes to or from the X chromosome. The co-adaptation hypothesis predicts that the co-transmission of X-linked and organellar genes should result in their co-adaptation, in which selection on beneficial epistatic interactions results in an over-representation of cyto-nuclear genes on the X chromosome relative to autosomes (Rand et al. 2004; Drown et al. 2012). In contrast, the sexual conflict hypothesis predicts the opposite chromosomal distribution, with more cyto-nuclear genes occurring on autosomes to alleviate mutation load in males. To date, empirical evidence for these hypotheses are mixed. Drown et al. (2012) used previously published reference genomes to examine the chromosomal distribution of N-mt genes in 16 vertebrates and found a strong under-representation of such genes on the X chromosomes relative to autosomes in 14 mammal species, but not in two avian species with ZW sex determining systems; note that the co-adaptation hypothesis does not predict that ZW systems should show a bias in the distribution of cyto-nuclear genes. Dean et al. (2014) included seven additional species in their analysis with independently derived sex chromosomes and found that the under-representation of N-mt genes on the X chromosome was restricted to therian mammals and *Caenorhabditis elegans*.

Here, we use sex-linked and autosomal transcriptome sequences to investigate the chromosomal distributions of cyto-nuclear interactions in the dioecious annual plant *Rumex hastatulus* (Polygonaceae). Examining cyto-nuclear interactions within a plant species is of interest for several reasons (see Sloan 2014). First, plants carry an additional maternally inherited organellar genome that is absent in animals, the chloroplast genome. This provides an opportunity to compare the chromosomal distribution of two independent kinds of cyto-nuclear interacting genes: nuclear-mitochondrial and nuclear-chloroplast. Second, whereas animal sex chromosomes evolved hundreds of millions of years ago (180 MYA in mammals and 140 MYA in birds; Cortez et al. 2014), the origin of plant sex chromosomes is a more recent event (Charlesworth 2013). In *R. hastatulus,* sex chromosomes are thought to have evolved approximately 15-16 MYA (Navajas-Perez et al. 2005) and genes on the Y chromosome show evidence of degeneration, resulting in a considerable proportion of genes that are hemizygous on the X chromosome (Hough et al. 2014). *Rumex hastatulus* therefore provides an opportunity to test whether the early changes involved in sex chromosome evolution are associated with a concomitant shift in the chromosomal location of N-mt or N-cp genes. Moreover, the presence in this system of X-linked genes that have recently become hemizygous provides an opportunity to compare the chromosomal distributions of X-linked genes that are hemizygous versus those that have retained Y-linked alleles (X/Y genes). Hemizygous genes are particularly good candidates for evaluating evidence for co-adaptation and/or sexual conflict because of their relatively older age (Hough et al. 2014), and because beneficial mutations in such genes are exposed to positive selection regardless of dominance and may therefore spread more rapidly.

## Methods

### Gene identification and functional annotation

We used sex-linked and autosomal transcriptome sequence data for *R. hastatulus* reported in Hough et al. (2014; GenBank Sequence Read Archive accession no. SRP041588), and obtained three sets of genes with which to test for an over- or under-representation of nuclear-mitochondrial or nuclear-chloroplast genes. In total our analyses included 1167 autosomal genes, 624 X-linked genes, and 107 hemizygous X-linked genes. The X-linked and hemizygous X-linked genes were shared between sex chromosome systems in this species (see Hough et al. 2014; *Methods* and *SI Appendix* for full details regarding the identification of such genes from transcriptome sequence data). For autosomal genes, we included those previously identified as confidently autosomal in both *R. hastatulus* sex chromosome systems, as well as those uniquely identified in the XYY system. For each gene set, we queried the sequences translated in all reading frames against the *A. thaliana* protein database using the BLASTx homology search implemented in Blast2GO (Conesa et al. 2005), with a significance threshold (BLAST ExpectValue) of 1 × 10^-3^, above which matches were not reported. We limited our searches to the *A. thaliana* protein database because sequence matches to this database returned more detailed functional information than is available for most other species in the NCBI plant database. We obtained BLASTx results for 1073 autosomal genes (90%), 567 X-linked genes (90%), and 95 hemizygous genes (89%). Gene Ontology (GO) terms associated with the hits from BLASTx queries were then retrieved using the ‘Mapping’ function in Blast2GO, which used BLAST accessions to link the queried sequences to functional information stored in the GO database (The Gene Ontology Consortium 2008). Gene names were retrieved using NCBI mapping files ‘gene info’ and ‘gene2accession’, and GO terms were assigned to query sequences using the ‘Annotation’ function with an E-Value-Hit-Filter of 1 × 10^-6^ and an annotation cut off of 55 (default parameters). Finally, we ran InterProScan (Quevillon et al. 2005) to retrieve sequence domain/motif information and merged the corresponding annotations with previously identified GO terms. This procedure generated output files containing GO ID’s and functional descriptions for each gene in our data set (files will be uploaded to GitHub). The numbers of genes in our final data set with functional annotations and N-mt and N-cp GO annotations are summarized in Table 1.

**Table 1.** Number of genes in our data set, including those for which we obtained functional annotations (see Methods) and those with nuclear-mitochondrial and nuclear-chloroplast GO annotations.

| Gene sets | Autosomal | X-linked | X-hemizygous |
| --- | --- | --- | --- |
| Original data set | 1167 | 624 | 107 |
| With annotation | 1073 | 567 | 95 |
| With N-mt GO ID | 194 | 94 | 13 |
| With N-cp GO ID | 222 | 102 | 22 |

### Statistical analyses

We used a similar approach to Drown et al. (2012) and Dean et al. (2014) and estimated the number of N-mt and N-cp genes on the X chromosome and autosomes, and then compared each of these estimates to an expected number. The expected number of N-mt genes was obtained by calculating the product of the proportion of all genes in the data set with mitochondrial annotations (matching GO:0005739) and the number of annotated genes in a given gene set. The expected numbers of N-cp genes were calculated similarly, using GO:0009507. We then calculated the ratios of the observed-to-expected numbers for both N-mt and N-cp genes in each gene set. The observed-to-expected ratio is expected to equal one when there is no under- or over-representation, and greater than one when there is an over-representation. We note that, unlike for X-linked genes, we did not have information regarding the particular chromosome locations for autosomal genes, and therefore could not obtain the expected numbers of N-mt and N-cp genes per-autosome as in previous studies (Drown et al. 2012; Dean et al. 2014). The expected numbers were thus calculated assuming that the set of autosomal genes represented a random sample of the autosomal chromosomes in this species, which is likely a valid assumption given that the sequences were obtained using whole transcriptome shotgun sequencing (Hough et al. 2014). Calculating the expected-to-observed ratios across X-linked, autosomal, and X-hemizygous genes thus allowed us to determine whether any of these gene sets contained an under- or over-representation of N-mt and N-cp genes compared to the expectation based on the proportion of such genes in the full data set. We tested the significance of over- or under-representation using Fisher’s exact tests, and calculated 95% confidence intervals for the numbers of N-mt or N-cp genes using 10,000 replicate bootstrapped samples. Given our sample sizes of genes with annotations (Table 1), Fisher’s Exact Tests allowed us to test for differences in the proportions of cyto-nuclear genes on autosomes versus the X-chromosome that were on the order of 5% with ∼80% power, whereas power was reduced for smaller differences (Supplementary Material). Similarly, for hemizygous X-linked genes, we calculate that differences of approximately∼10% could be detected with ∼80% power. All data analysis was done in R (R Development Core Team 2013; scripts will be available for download from GitHub).

## Results and Discussion

It has been suggested that cyto-nuclear genes may be either over- or under-represented on the X chromosome compared to autosomes, depending on whether their interactions are driven by co-adaptation or sexual conflict (Rand et al. 2001; Drown et al. 2012; Hill and Johnson 2013; Dean et al. 2014; Rogell et al. 2014). We annotated sex-linked and autosomal transcriptome sequences to test these predictions in the dioecious plant *R. hastatulus*. We found that neither mitochondria- or chloroplast-interacting nuclear genes were under- or over-represented on the X chromosome (Fisher’s exact test, *P* = 0.4947 and *P* = 0.3074, respectively; Figure 1). This pattern indicates that neither the co-adaptation nor the sexual conflict hypothesis alone is sufficient to explain the chromosomal distribution of cyto-nuclear genes in *R. hastatulus*.

**Figure 1.**
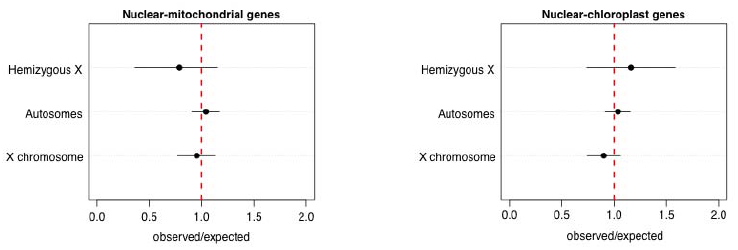
Representation of the chromosomal location of cyto-nuclear genes in *Rumex hastatulus*. Dots represent the observed to expected ratio of mito-nuclear (N-mt) and chloro-nuclear (N-cp) genes on autosomes, the X chromosome, and hemizygous X genes, with the 95% confidence intervals estimated by bootstrapping (10,000 replicates). The vertical dotted line at 1 represents no over- or under-representation.

There are several factors that are expected to be important in determining cyto-nuclear gene distributions, and these may explain the lack of bias in *R. hastatulus*. For example, under both the co-adaptation and sexual conflict hypotheses, the age of the sex chromosomes will determine the extent to which selection (either for co-adaptation, or sexual antagonism) has had time to operate, which depends on the rate of gene movement onto and off of the sex chromosomes. Whereas previous studies of cyto-nuclear genes in animals have focused almost exclusively on ancient sex chromosome systems (Drown et al. 2012; Dean et al. 2014; Rogell et al. 2014), our study focused on a dioecious plant species in which sex chromosomes evolved more recently (∼15 MYA; Navajas-Perez et al. 2005), and many genes likely stopped recombining much more recently (Hough et al., 2014). The lack of bias in the chromosomal distribution of cyto-nuclear genes may therefore reflect the recent time scale of sex chromosome evolution rather than the absence of biased gene movement. The relatively young age of sex chromosomes may also have played a role in the lack of bias reported in the sex and neo-sex chromosomes in three-spined stickleback, which evolved ∼10 MYA (Kondo et al. 2004) and ∼2 MYA, respectively (Natri et al. 2013). Comparative studies of sex chromosomes of different age will be central for understanding the rate at which organellar gene movement occurs.

In addition to being evolutionarily older, X-linked hemizygous genes are expected to show a greater effect of over-or under-representation than genes with both X- and Y-alleles because recessive mutations (involved in either co-adaptation or sexual conflict) will be exposed to selection instead of masked by an alternate allele in a heterozygous genotype. We detected a slightly greater under-representation of X-hemizygous N-mt genes compared to autosomes or X-genes with retained Y-alleles, but the effect was not statistically significant (*P* = 0.4947). The opposite pattern was evident for N-cp genes, which were slightly over-represented on hemizygous genes, but again this effect was not significant (*P* = 0.3074). A larger sample of hemizygous genes would be required to more confidently assess whether such genes are in fact more often involved in cyto-nuclear interactions than other genes on the X chromosome, and to test whether the opposite pattern for N-mt and N-cp hemizygous genes is a result of a different rate of nuclear gene transfer between mitochondrial and chloroplast genomes. In particular, the smaller number of hemizygous X-linked genes in our data set implies that power was reduced for this comparison, such that a ∼5% difference could only be detected with ∼60% power (see Supplementary Material).

Another factor that will affect the chromosomal distribution of cyto-nuclear genes is the number of N-mt and N-cp genes that were located on the autosome from which the sex chromosomes evolved. Since the origins of mitochondria and chloroplasts both vastly predate that of sex chromosomes (1.5-2 BYA compared to < 200 MYA; Dyall et al. 2004; Timmis et al. 2004; Cortez et al. 2014), gene transfer from organellar genomes to the nuclear genome began long before the evolution of sex chromosomes. A bias in the chromosomal distribution of cyto-nuclear genes in either direction may therefore arise if the ancestral autosome was particularly rich or poor in cyto-nuclear genes. Indeed, it is striking that autosomes in the animal species previously examined exhibited extensive variation in the relative number of N-mt genes (see Drown et al. 2012 Figure 1 and Dean et al. 2014 Figure 1 and Figure 2). That the ancestral number of N-mt and N-cp genes is likely to be important is highlighted by the fact that the majority of genes involved in mitochondrial DNA and RNA metabolism in *A. thaliana* are found on chromosome III (Elo et al. 2003). If such a biased autosomal distribution of organellar variation is representative of the ancestral sex chromosomes, the X chromosome could carry significantly more N-mt or N-cp genes because of this ancestral gene number rather than a biased rate of gene movement. This effect is likely exacerbated in early sex chromosome systems, where the majority of genes may not have experienced opportunities for movement. Genetic mapping and comparative genomic studies of genes that have transferred from organellar genomes after the origin of sex chromosomes may provide a means to control for ancestral differences in gene number and provide a better test of biases in organellar-nuclear gene movement.

To conclude, our study is the first investigation of the extent to which co-adaptation and sexual conflict have shaped the chromosomal distribution of cyto-nuclear genes in a plant species with sex chromosomes. We found no sign of under- or over-representation of either N-mt or N-cp genes on the X chromosome, implying that neither co-adaptation nor sexual conflict alone can explain the chromosomal distributions of these genes. Instead, we suggest that additional factors, including the age of sex chromosomes and the time that has elapsed since X-Y recombination became suppressed, are likely to have been important determinants of the patterns we observed. To determine whether the lack of under-representation of mito-nuclear genes on the X chromosome reflects an absence of gene movement, future studies should focus on quantifying rates of gene movement after sex chromosome origination, and consider the extent to which neutral processes including the number of mito-nuclear genes on ancestral sex chromosomes have played an important role in shaping the current chromosomal distributions of such genes. Cyto-nuclear conflict and co-evolution have undoubtedly played a major role in many aspects of genome evolution in both plant and animal systems, and the previously reported evidence from therian mammals and *C. elegans* (Drown et al. 2012; Dean et al. 2014) suggests that sexual conflict and co-adaptation might represent important mechanisms driving chromosomal gene movement; however, it remains unclear whether these processes have also shaped the chromosomal distribution cyto-nuclear genes in plants.

## Acknowledgements

We thank Rebecca Dean and Devin M Drown for comments on the manuscript. This research was supported by Discovery Grants to SCHB and SIW from the Natural Sciences and Engineering Council of Canada. JH was supported by an Ontario Graduate Fellowship and JAÅ by a Junior Fellowship from Massey College.

